# Electrophysiological indices of distractor processing in visual search are shaped by target expectations

**DOI:** 10.1101/2022.12.21.521409

**Authors:** Dirk van Moorselaar, Changrun Huang, Jan Theeuwes

## Abstract

Although in many cases salient stimuli capture attention involuntarily, it has been proposed recently that under certain conditions the bottom-up signal generated by such stimuli can be proactively suppressed. In support of this signal suppression hypothesis, ERP studies have demonstrated that salient stimuli that do not capture attention elicit a distractor positivity (P_D),_ a putative neural index of suppression. At the same time, it is becoming increasingly clear that regularities across preceding search episodes have a large influence on attentional selection. Yet to date, studies in support of the signal suppression hypothesis have largely ignored the role of selection history on the processing of distractors. The current study addressed this issue by examining how electrophysiological markers of attentional selection (N2pc) and suppression (P_D_) elicited by targets and distractors respectively were modulated when the search target randomly varied instead of being fixed across trials. Results showed that while target selection was unaffected by this manipulation, both in terms of manual response times, as well as in terms of the N2pc component, the P_D_ component was reliably attenuated when the target features varied randomly across trials. This result demonstrates that the distractor P_D_, which is typically considered the marker of selective distractor processing cannot unequivocally be attributed to suppression only, as it also, at least in part, reflects the upweighting of target features.

## Introduction

The understanding that attentional selection is not only determined by the interaction between top-down and bottom-up processes, but is also strongly influenced by previous selection episodes has revived the attentional capture debate (Luck, Gaspelin, Folk, Remington, & Theeuwes, 2021; van Moorselaar & Slagter, 2020). This debate is centred around the perplexing puzzle that on the one hand salient, yet irrelevant stimuli often appear to involuntarily capture attention, while at the same time such involuntary capture would make simple everyday tasks nearly impossible. Recent research demonstrating that learning about regularities in the environment does not only prioritize target properties, such as the location and features of the target (Chun & Jiang, 1998), but can also result in the suppression of task irrelevant information (Ferrante et al., 2018; Leber, Gwinn, Hong, & O’Toole, 2016; Sauter, Liesefeld, & Müller, 2019; van Moorselaar & Theeuwes, 2022; Wang & Theeuwes, 2018), may provide a missing piece to this puzzle.

As an initial resolution, the signal suppression hypothesis (Gaspelin & Luck, 2018b; Sawaki & Luck, 2010) was put forward, which elegantly incorporates the seemingly conflicting sides of the attentional capture debate. Specifically, the signal suppression hypothesis posits that while salient stimuli all generate a bottom-up “attend-to-me” signal and hence compete for attention, this signal can be proactively suppressed if the attentional system is appropriately configured (Luck et al., 2021). This idea is supported by empirical findings that salient stimuli automatically capture attention when the target is defined as a unique element in the display and therefore there is no clear search goal (e.g., the target is defined as the unique shape in the search display; singleton detection mode). By contrast, when the target is a specific shape embedded in a display of heterogeneous shapes and observers hence engage in what is called feature search mode (Bacon & Egeth, 1994), allowing them to impose top-down control, which not only eliminates attentional capture (Folk, Remington, & Johnston, 1992), but under certain conditions can even result in below baseline suppression (Gaspelin & Luck, 2019). While support for this hybrid model of attentional capture has accumulated through converging evidence of psychophysics (e.g.,)(Adam, Patel, Rangan, & Serences, 2021; Gaspelin, Leonard, & Luck, 2015), and eye movement studies (e.g.,)(Gaspelin, Gaspar, & Luck, 2019; Gaspelin, Leonard, & Luck, 2017), some influential studies relied on electrophysiological indices related to attentional selection (the N2pc) and suppression (the P_D_) (Eimer, 1996; Luck & Hillyard, 1994a, 1994b) and suppression (Hickey, Di Lollo, & McDonald, 2009).

Although the N2pc, which is a negative going deflection occurring around 200-300 msec after stimulus presentation that is larger over the hemisphere contralateral to the attended location, is unequivocally considered as an index of covert attention (Luck, 2012; Woodman & Luck, 2003), the Pd is less well characterized. Nevertheless, the P_D,_ which is much like the inverse of the N2pc, as it presents itself as a larger contralateral positive deflection to a to-be-ignored rather than a task-relevant stimulus, is commonly presented as a putative neural index of suppression (Gaspelin & Luck, 2018b; Hickey et al., 2009) and has become a tool to study whether salient distractors can be suppressed proactively (Drisdelle & Eimer, 2021; Stilwell, Egeth, & Gaspelin, 2022; van Moorselaar, Daneshtalab, & Slagter, 2021; van Moorselaar, Lampers, Cordesius, & Slagter, 2020; van Moorselaar & Slagter, 2019; Wang, van Driel, Ort, & Theeuwes, 2019). Indeed, various strands of evidence link the P_D_ to a suppressive mechanism; the Pd often appears exclusively in response to distractors in the absence of a N2pc (e.g.,)(Gaspar & McDonald, 2014; Jannati, Gaspar, & McDonald, 2013; Sawaki & Luck, 2010), it has a larger amplitude on a subset of trials with the fastest responses (McDonald, Green, Jannati, & Di Lollo, 2013) and is no longer found when observers fail to avoid an eye-movement towards the distractor (Weaver, van Zoest, & Hickey, 2017). However, arguably the most convincing evidence thus far linking the P_D_ to suppression comes from a study by Gaspelin and Luck (2018a) that observed a correlation between the magnitude of below baseline behavioural suppression and the magnitude of the P_D_ component (see also)(Feldmann-Wüstefeld, Brandhofer, & Schubö, 2016; Gaspar & McDonald, 2014).

There is thus good reason to believe that in many situations the P_D_ can be linked to a suppressive mechanism (van Moorselaar & Slagter, 2020). At the same time it should be noted that studies examining below baseline suppression through feature search by means of the P_D_, thus far have largely ignored the potential modulation by regularities across search displays (Theeuwes, Bogaerts, & van Moorselaar, 2022; van Moorselaar & Slagter, 2020). In this respect, it is noteworthy that in the work by Gaspelin and colleagues (Gaspelin & Luck, 2018a; Stilwell et al., 2022) examining proactive distractor suppression via the P_D_ not only the distractor feature (i.e., its colour) but also the target features (i.e., both its colour and shape) were fixed across trials. In other words, the experimental designs used by Gaspelin and colleagues strongly induced, be it via implicit learning (Theeuwes et al., 2022) or an explicit top-down process (Wolfe, 1994), predictions regarding both targets and distractors. It is therefore unclear to what extent the observed electrophysiological markers of distractor processing reflect pure distractor-feature suppression, as typically assumed, or alternatively, at least partly, also reflect target-feature upweighting (Chang & Egeth, 2019).

While the P_D_ is exclusively elicited by to-be-ignored items, which seems to favour an interpretation in terms of suppression, there is reason to believe that it may in part also reflect upweighting of target features. As also acknowledged by updated versions of the signal suppression hypothesis (Luck et al., 2021), distractor inhibition, often results not only from proactive suppression of static and hence predictable distractor features (Vatterott & Vecera, 2012), but in part also reflects upweighting of predictable target features. For example, by modifying the capture-probe technique, a technique used to read out attentional processing across search displays, such that it could independently dissociate between distractor suppression and target feature upweighting, Chang and Egeth (2019, 2021) demonstrated that both distractor suppression and target feature upweighting contributed to distractor inhibition. Moreover, based on a more systematic manipulation of colour similarity between the target and neutral filler items in the display, Oxner and colleagues (2022) even concluded that apparent proactive distractor suppression could be entirely explained by global target feature enhancement. While future work is necessary to further understand the interplay between suppressive and upweighting mechanisms in the context of distractor suppression, to the very least these findings illustrate the importance of disambiguating target and distractor effects in P_D_ research. Given that in recent years the P_D_ and its modulation has been widely used by both sides of the attentional capture debate, it is critical to establish to what extent it represents pure distractor processing independent from learned expectations regarding task relevant features, which to the best of our knowledge has not yet been done experimentally.

To examine the extent to which, if at all, the P_D_ component also reflects upweighting of target features, we adopted the paradigm used by Gaspelin and colleagues that reliably produces below baseline suppression and a robust distractor P_D_ (Gaspelin & Luck, 2018a; Stilwell et al., 2022), but varied whether or not target features were static across trials. Specifically, while in half of the experiment both the target and the distractor feature were fixed across trials (as in previous studies), in the other half of the experiment, target features (i.e., shape and colour) varied randomly across trials (see Figure 1A). This design allowed us to establish whether the P_D_ as observed in previous studies reflects pure distractor suppression, or alternatively, at least partly, is driven by upweighting of static target features. Critically, if the P_D_ purely reflects distractor suppression, under the current conditions the P_D_ component should not be modulated by cross-trial variation of the target features.

**Figure 1.**
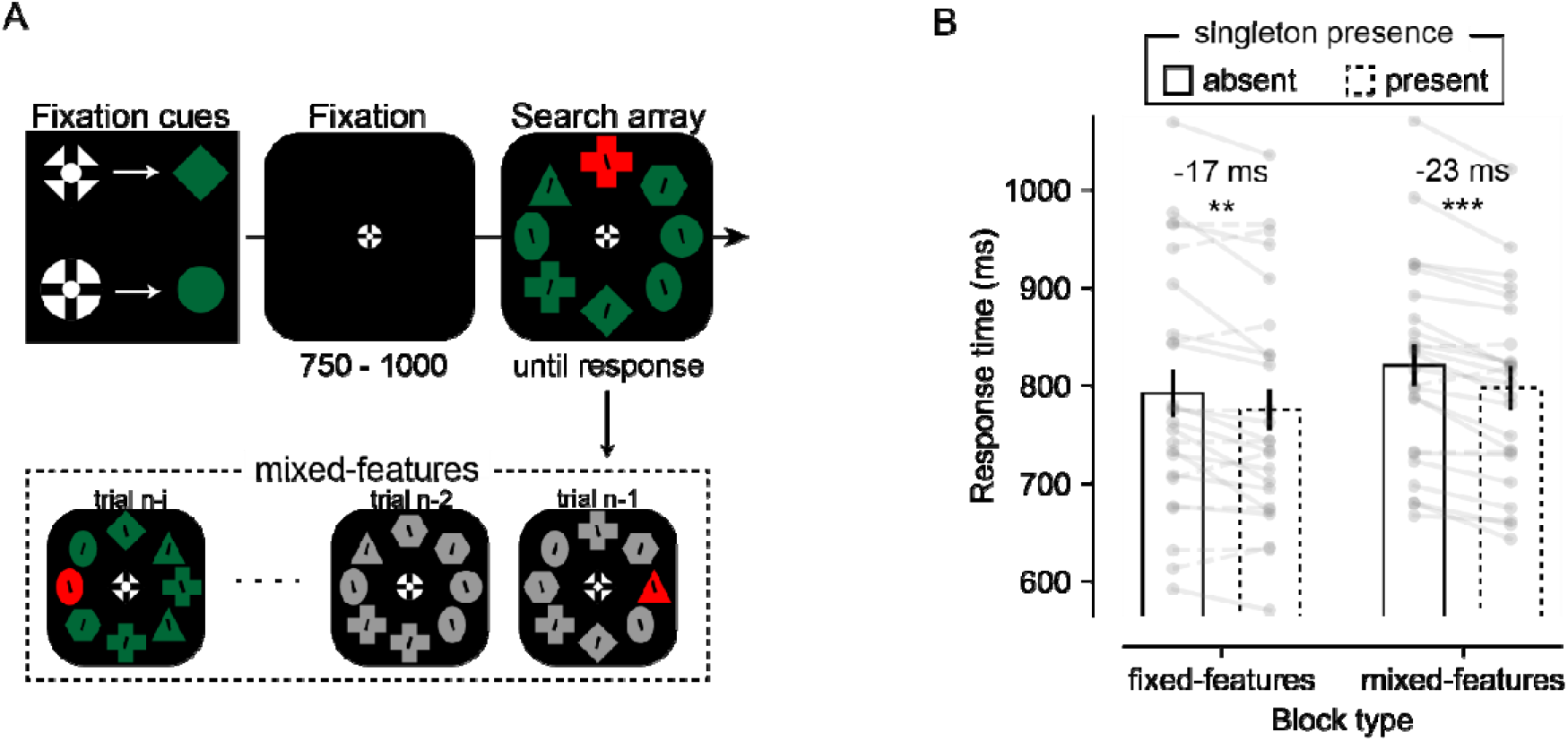
Schematic of the experimental procedure and behavioural results. **A)** On each trial, a heterogeneous set of shapes was presented in a circular configuration around fixation. The target shape (i.e., diamond or circle) was cued by the shape of the fixation marker (see top left of the figure). In singleton present displays, one of the shapes had a unique colour (red or green; counterbalanced across participants) which was fixed throughout the entire experiment. Participants (n = 24) were instructed to report the orientation of the line inside the shape cued by the fixation marker (i.e., diamond or circle). In the fixed-features condition (half of the experiment) this target shape, as well as the target colour was static across trials. By contrast, in the mixed-features condition, both the target shape (diamond or circle) as well as the target colour (red/green or grey) varied randomly across trials. Search stimuli were taken from the Stilwell et al. (2022) study on which the current study was based. **B)** The singleton presence benefit was not modulated by target regularities: In the fixed-features condition participants detected the target faster on singleton present (dashed bar) versus singleton absent (solid bar) displays (*m* _absent_ = 792.0; *m* _present_ = 775.2; ∆*m = 16*.*8*.; *n = 24;* two-tailed *p* = 0.006; *d* = 0.61; 95% CI = 5.3 – 28.4). In the mixed-features condition, the same benefit was observed (*m* _absent_ = 820.4; *m* _present_ = 797.2; ∆*m = 23*.*2*.; *n = 24;* two-tailed *p* < 0.001; *d* = 1.2; 95% CI = 15.3 – 31.1) resulting in a unreliable Block type by Singleton presence interaction (*F* (1, 23) = 1.3, *p* = 0.27, = 0.052; BFexcl = 3.1). The height of each bar reflects the population average, and error bars represent 95% within-subject confidence intervals (Morey, 2008). Data from each participant are represented as grey dots, connected by solid lines (i.e., singleton presence benefit) or dashed lines (i.e., attentional capture).

### Open practice statement

Experiment and analyses were based on the *OSF Preregistration template* at Open Science Framework (https://osf.io/9827w/). Analyses that diverge from the preregistration are described as exploratory. Deidentified data for all experiments along with the data-analysis scripts (custom Python 3 scripts) will be posted alongside the preregistration upon publication. All code for running the experiment will also be made available here.

## Methods

### Participants

A planned number of 24 participants (mean age = 21 years, range = 19 - 26; 21 female), participated in the experiment, in exchange for course credit or monetary compensation (10 euros per hour). Sample size was based on our previous work (van Moorselaar et al., 2021) and previous studies on which the current study was based (Gaspelin & Luck, 2018a; Stilwell et al., 2022). Eight participants were replaced because they failed to maintain fixation during the window of interest in a large subset of trials (*N* = 6), because automatic artefact rejection resulted in removal of too many trials (*N* = 1), or because their accuracy deviated more than 2.5 *S*.*D*. from the group mean (*N* = 1). All participants gave their informed consent prior to the start of the study, which was approved by the Ethical Review Committee of the Faculty of Behavioural and Movement Sciences of the Vrije Universiteit Amsterdam.

### Apparatus, material and procedure

The experiment, which took place in a dimly lit room on a 23.8-inch ASUS ROG STRIX XG248 LED monitor with a 240 Hz refresh rate, was created using OpenSesame (Mathôt, Schreij, & Theeuwes, 2012) utilizing Psychopy functionality (Peirce, 2009). Participants were positioned 60 cm away from the screen using a desk mounted chinrest. The eyes were tracked on- and offline using an Eyelink 1000 (SR research) eye tracker tracking the left eye with a 1000 Hz sampling frequency (one subject had a sampling frequency of 2000 Hz) and participants heard a beep each time fixation was broken by more than 2° of visual angle. At the start of the experiment the eyes were calibrated via a five dots calibration procedure until spatial error for each dot position was smaller than 1 degree of visual angle. Drift correction was applied every 80 trials (i.e., at the start and halfway a block), when deemed necessary the calibration procedure was repeated. EEG data were recorded at a sampling rate of 512 Hz with default settings using a 64-electrode cap with electrodes placed according to the 10-10 system (Biosemi ActiveTwo system; biosemi.com). Vertical and horizontal EOG (VEOG/HEOG) were recorded via external electrodes placed ∼2 cm above and below the eye, and ∼1 cm lateral to the external canthi, respectively.

The paradigm was modeled after Stilwell et al. (2022). Each trial started with a randomly jittered black display (100 – 400 ms) followed by a randomly jittered fixation display (750 – 1000 ms). This display contained either a circular or a diamond shape with an embedded cross hair, a combination that has been shown to improve stable fixation (Thaler, Schütz, Goodale, & Gegenfurtner, 2013). Critically, the shape of the fixation marker signaled the target shape in the subsequent search display. Each search display contained eight shapes in a circular configuration (radius 3°) around the fixation marker, each with a black line tilted left or right (14° around the vertical plane) in their center. Individual shapes were selected from a stimulus pool of triangles (radius 0.7°), hexagons (1.2° by 1.2°), ovals (1.5° by 0.9°), crosses (1.2° by 1.2°), diamonds (1.3° by 1.3°) and circles (radius 0.6°). Selection was such that each display contained the target shape (i.e., diamond or circle) and seven shapes randomly selected from the remaining shapes in the stimulus pool with replacement, with the restriction that each individual non-target shape appeared two times at the most to ensure high display heterogeneity.

Individual shapes were either red (RGB: 253, 34, 34), green (RGB: 90, 174, 20) or grey (RGB: 146, 147, 153). In the fixed-feature condition both the shape (circle or diamond; counterbalanced across participants) and the colour of the target (red or green; counterbalanced across participants) were held constant. By contrast, in the mixed-feature condition both the target shape and target colour varied randomly across trials (counterbalanced across trials in a block) such that in half of the displays the target colour was grey, whereas in the other half the target colour matched the target colour in the fixed-feature condition. Whereas in 25% of trials all stimuli in the search display had the same colour (i.e., singleton absent displays), in the remaining trials (i.e., singleton present displays) one of the non-target shapes was rendered in a unique colour (red or green; counterbalanced across participants) that was held constant throughout the experiment. These singleton absent and present displays were not randomly intermixed, but instead singleton absent displays were grouped together at the start or at the end of an experimental block (alternating between blocks) such that any singleton presence benefit could not be attributed to a surprise induced by infrequent distractor absent displays. Across all display configurations targets and singleton distractors appeared with equal probability selectively at positions along the vertical and horizontal axis.

At the beginning of each session, it was made explicit that the distractor singleton was irrelevant to the task at hand and should thus be ignored. Participants were instructed to keep their eyes at fixation and covertly search for the shape that matched the fixation shape on the current trial and indicate the orientation of the line segment within this target shape via button press (i.e., ‘z’ or ‘/’ button) as quickly as possible while keeping the number of errors to a minimum. In case of an incorrect response or missing response a 200 Hz tone lasting 300 ms was played, which was accompanied by the text “too slow!” in case participants did not respond within 2000 ms.

The experiment consisted of 12 experimental blocks of 160 trials (6 consecutive blocks for each condition; order counterbalanced across participants) preceded by sequences of 32 training trials in the mixed-feature condition, which repeated until mean accuracy was above 70%. At the start of each new block participants were informed about the dynamics of the upcoming block (mixed-feature or fixed-feature condition) and whether distractor present/absent displays were grouped at the start or at the end of the upcoming block. Halfway each block there was a 15 sec mandatory break to rest the eyes. After each block participants received feedback on their performance (i.e., mean reaction time (RT) and accuracy).

### Behavioural analysis

All data were preprocessed in a Python environment (Python Software Foundation, https://www.python.org/). Analyses were limited to RT data of correct trials only. RTs were filtered in a two-step trimming procedure: trials with RTs shorter than 200 msec were excluded, after which data were trimmed based on a cutoff value of 2.5 SD from the mean per participant. Remaining RTs were analyzed with repeated measures ANOVAs with within-subject factors Block type (fixed-features, mixed-features) and Singleton presence (present, absent), followed by planned comparisons with paired *t*-tests using JASP software (<u>JASP-TEAM, 2018</u>). In case of insignificant interactions, we also report BF_excl_ which reflects the comparison between the interaction and equivalent models stripped of the effect.

### EEG preprocessing

EEG data, which was re-referenced offline to the average of the left and right earlobe, was first high-pass filtered using a zero-phase ‘firwin’ filter at .1 Hz to remove slow drifts. Continuous EEG was subsequently epoched from −700 – 1100 ms relative to search display onset (to avoid filter artefacts during automatic artefact rejection; see below). Prior to trial rejection, malfunctioning electrodes as identified during recording (*M* = 0.6, range = 0-2) were temporarily removed. As a first artifact removal step, ICA as implemented in MNE (method = ‘picard’) was performed on 1 Hz filtered epochs to remove eye-blink components selectively from the 0.1 Hz filtered data. Next noise contaminated epochs within a −200 – 600 ms were identified using an adapted version of an automatic trial-rejection procedure. To specifically capture muscle activity, the EEG signal was filtered using a 110 – 140 Hz band-pass filter and subsequently transformed into Z-scores. A subject specific Z-score threshold was then set based on within-subject variance of Z-scores (Vries et al., 2017). Moreover, to reduce the number of false alarms, rather than immediate removal of epochs exceeding the Z-score threshold, per marked epoch the five electrodes that contributed most to accumulated Z-score within the time period containing the marked artefact were identified. Then in an iterative procedure, the worst five electrodes per marked epoch were interpolated using spherical splines (Perrin et al., 1989) one by one, checking after each interpolation whether that epoch still exceeded the determined Z-score threshold. Epochs were selectively dropped when after this iterative interpolation procedure, the Z-score threshold was still exceeded. Finally, malfunctioning electrodes were interpolated using spherical splines (Perrin, Pernier, Bertrand, & Echallier, 1989).

### ERP analyses

ERP analyses were limited to trials without identified eye movements. For this purpose, trials with a fixation deviation >1° of visual angle correction in a segment of continuous data of at least 40ms in the time window −200 - 400-ms after drift correction using pre-stimulus data (van Moorselaar & Slagter, 2019) were excluded. We focused the analysis of distractor- and target-elicited ERP waveforms on electrodes PO7/8, which were chosen a priori based on previous studies examining the Pd (Gaspelin & Luck, 2018a; Stilwell et al., 2022). Epochs were baseline corrected using a −200 – 0-ms pre-stimulus baseline period. To enable isolation of lateralized distractor- and target-specific components, the analyses focused on trials in which the stimulus of interest (distractor or target) was presented to the left or right of fixation, while the other stimulus was presented on the vertical meridian or absent. Waveforms evoked by the various search displays were collapsed across left and right visual hemifield and left and right electrodes to produce separate waveforms for contralateral and ipsilateral scalp regions. Lateralized difference waveforms were then computed by subtracting the ipsilateral waveform from the corresponding contralateral waveform. Time windows of interest were centered around the positive peak (±55-ms) of the grand mean waveform (i.e., averaged across conditions) in distractor tuned analyses, and around the negative peak (±37.5-ms) of the grand mean waveform in target tuned analyses.

## Results

### Search times

Exclusion of incorrect responses (7.5%) and data trimming (2.9%) resulted in an overall loss of 10.5% of behavioural data. Figure 1A depicts the mean RTs for singleton present and absent displays in the two search conditions. Efficiency of target selection appeared to not differ as a function of whether target features were static or varied randomly across trials (*F* (1, 23) = 2.57, *p* = 0.12, 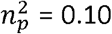; BFexcl = 0.24). Critically, there was a clear benefit when the search array contained a singleton distractor (main effect Singleton presence: *F* (1, 23) = 26.84, *p* < 0.001, 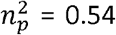), independent of whether the target features (i.e., shape and colour) were fixed or varied randomly across trials (interaction Singleton presence and Block type: *F* (1, 23) = 1.26, *p* = 0.27, 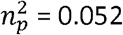; Bfexcl = 3.1). Replicating previous studies using heterogeneous search displays (Chang & Egeth, 2019; Gaspelin & Luck, 2018a; Stilwell et al., 2022), in the fixed-feature condition there was a reliable ∼17 msec singleton-presence benefit (*t* (23) = 3.01, *p* = 0.006, *d* = 0.61). This singleton-presence benefit was not reduced, but, if anything, larger (∼23 msec) and more reliable (*t* (23) = 6.05, *p* < 0.001, *d* = 1.24) when target features varied randomly across trials and hence the trial structure provided less opportunity to upweight relevant features for the upcoming search. Error rates did not reliably differ between singleton present and singleton absent trials, neither in fixed-features nor in mixed-feature conditions (interaction Search condition and Block type: *F* (1, 23) = 0.038, *p* = 0.85, 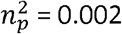; BFexcl = 3.56).

To further explore the effect of randomly switching target features, in the mixed-features condition we separated trials in which none of the target features, the target color, the target shape, or both target features repeated from one trial to the next. This analysis yielded no evidence that the singleton presence benefit was modulated by intertrial target feature priming (interaction Singleton presence and Target Feature repetition: *F* (3, 69) = 0.95, *p* = 0.42, 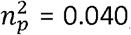; BFexcl = 5.17). Altogether, the behavioral results indicate that selection of the target, as well as ignoring the static singleton distractor color remained equally effective when target features varied randomly across trials compared to a condition with static target features. Given the putative roles of the N2pc, reflecting covert attentional selection (Luck, 2012; Woodman & Luck, 2003), and the Pd, reflecting suppression (Gaspelin & Luck, 2018a, 2018b), we should thus also not expect differences in the N2pc elicited by targets and, of especially interest here, the Pd elicited by distractors between mixed- and fixed-feature conditions.

### Electrophysiological results

We first examined the waveforms elicited by lateralized targets to characterize attentional selection across conditions. Previous studies examining target selection in heterogeneous search displays identified no reliable differences between difference waveforms elicited by lateral targets either with or without singleton distractors (Gaspelin & Luck, 2018a; Stilwell et al., 2022), as was also the case here. Therefore, in target tuned analysis (see Figure 2), we collapsed across singleton absent trials and trials with a distractor on the vertical midline (individual waveforms for distractor absent and distractor present displays are shown in supplementary Figures 1 and 2). As expected, lateralized targets elicited a more negative-going deflection in the contralateral compared to the ipsilateral waveform beginning at approximately 200-ms after search display onset. In line with behavior, this N2pc component appeared with approximately the same amplitude and time course when the target features were fixed across trials (i.e., fixed-features condition), and when they varied randomly across trials (i.e., mixed-features condition).

**Figure 2.**
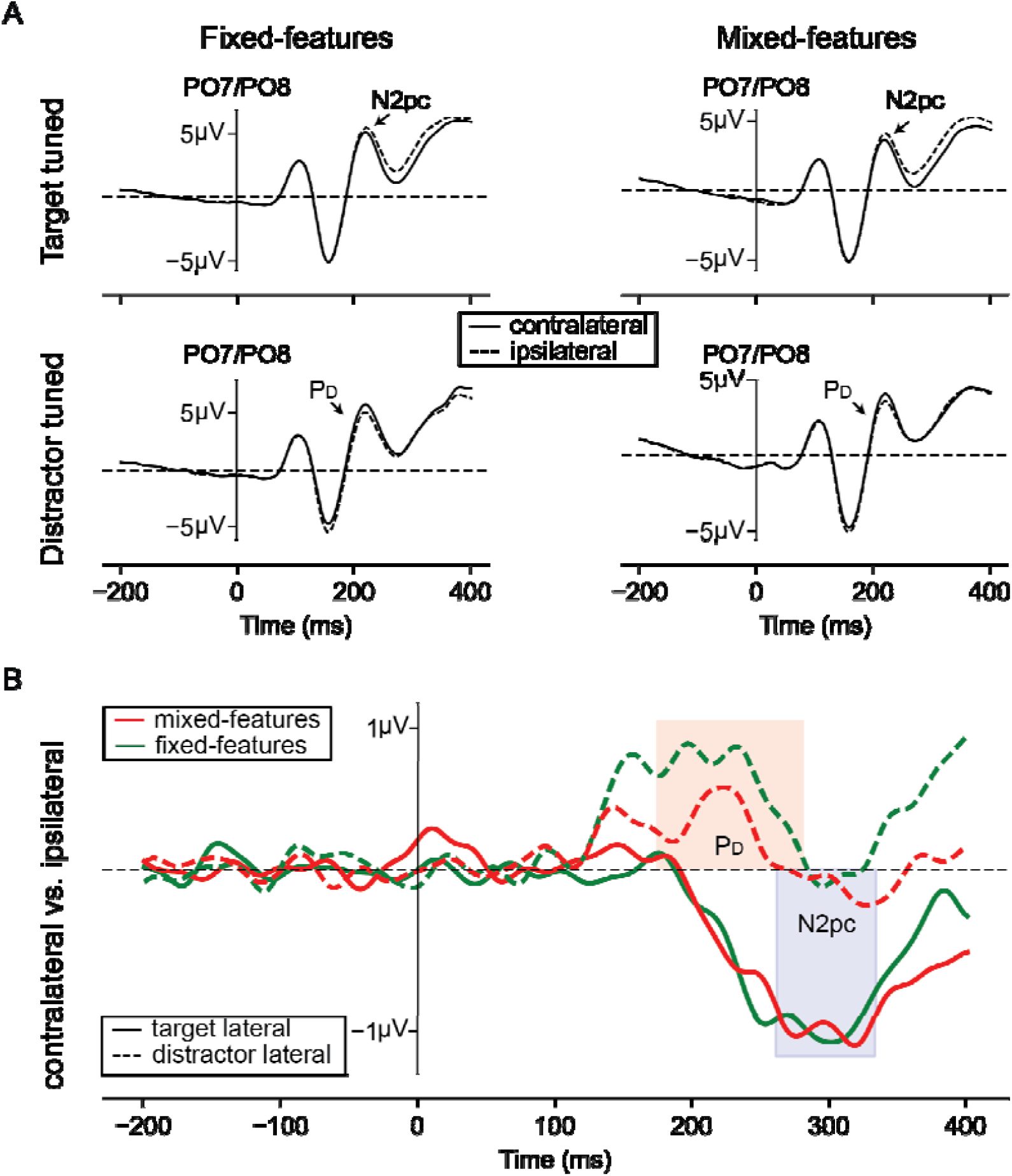
The distractor Pd, but not the target N2pc was modulated by target feature regularities. **A)** Electrophysiological results from search trials with lateral targets (target tuned ERPs; row 1) and with lateral distractors (distractor tuned ERPS; row 2), separately for fixed-features (column 1) and mixed-features (column 2) conditions, respectively. Ipsilateral (dashed lines) and contralateral (solid lines) waveforms reflect activity at electrode sides PO7/8. **B)** Difference waveforms between contra- and ipsilateral waveforms for target tuned (solid lines) and distractor tuned analyses (dashed lines), separately for fixed-features (green) and mixed-features (red) conditions. Shaded areas reflect the time windows of interest for the N2pc (blue) and the Pd (red) analyses). The N2pc elicited by lateral targets did not differ between the fixed-features condition and the mixed-features condition (*m* _fixed_ = −0.97µV; *m* _mixed_ = −1.0µV; ∆*m = 0*.*03*.; *n = 24;* two-tailed *p* = 0.89; *d* = 0.028; 95% CI = −0.37 – 0.43). In contrast, the Pd elicited by distractors was reliably attenuated, but nevertheless reliable, when target features varied randomly relative to a condition with static target features (*m* _fixed_ = 0.57µV; *m* _mixed_ = 0.25µV; ∆*m = 0*.*32*.; *n = 24;* two-tailed *p* = 0.031; *d* = 0.47; 95% CI = 0.04 – 0.89).

The N2pc components were measured as the mean amplitudes from 261 – 336-ms post stimulus and subsequently analyzed using a repeated measures ANOVA with within subjects’ factors Block type (mixed-features, fixed-features) and Hemifield (contralateral to target, ipsilateral to target). This analysis confirmed that the while the N2pc was reliable across conditions (main effect Hemifield: *F* (1, 23) = 28.58, *p* < 0.001, 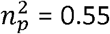), there was no evidence that it differed between block types (interaction Block type and Hemifield: *F* (1, 23) = 0.019, *p* = 0.89, 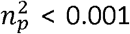; BFexcl = 3.91). Indeed, planned pairwise comparisons comparing contralateral vs. ipsilateral waveforms yielded reliable differences both in the fixed- (*t* (23) = 4.27, *p* < 0.001, *d* = 0.87) and the mixed-features (*t* (23) = 4.97, *p* < 0.001, *d* = 1.01) conditions.

While the target tuned analysis, mimicking the observed behavior, thus did not identify marked differences between conditions, somewhat surprisingly waveforms elicited by distractors appeared to be modulated by target feature regularities (see Figure 2). Like previous studies (Drisdelle & Eimer, 2021; Gaspelin & Luck, 2018a; Sawaki & Luck, 2010; Stilwell et al., 2022), the positive difference elicited by lateral distractors, corresponding to the Pd, had an earlier onset than the N2pc elicited by targets, seemingly consistent with the idea that the Pd signals proactive suppression (Gaspar & McDonald, 2014; Jannati et al., 2013; Sawaki & Luck, 2010). Yet, at odds with this idea, the Pd appeared to be attenuated when target features were no longer fixed across trials, suggesting that processing of singleton distractor as signaled by the Pd, at least to some extent, also reflects target feature upweighting rather than a pure suppressive process.

The Pd components were measured as the mean amplitudes from 174 to 284-ms post stimulus onset and subsequently analyzed in the same way as the N2pc components. Replicating previous studies there was a reliable lateralized positivity elicited by singleton distractors (main effect Hemifield: *F* (1, 23) = 11.60, *p* = 0.002, 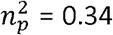). Critically, while planned pairwise comparisons showed that this Pd component was evident in both the fixed- (0.5 µV; *t* (23) = 3.38, *p* = 0.003, *d* = 0.69) and the mixed-features condition (0.2 µV; *t* (23) = 2.49, *p* = 0.022, *d* = 0.50), a Block type by Hemifield interaction confirmed that the Pd was reliably attenuated when target features were not static, but instead varied randomly across trials (*F* (1, 23) = 5.29, *p* = 0.031, 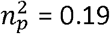)^1^. These findings demonstrate that when observers could not predict the target color in advance and were not consistently searching for the same shape, the P_D_ elicited by singleton distractors was reduced, but nevertheless reliable.

### Exploratory analyses

Visual inspection of Figure 2 suggests that the observed modulation of the P_D_ was especially apparent in the early time window of the Pd. Given that distractor positivity’s often contain an early and a late component (Feldmann-Wüstefeld & Vogel, 2019; van Moorselaar et al., 2021; Weaver et al., 2017), which have been speculatively linked to different cognitive processes, we explored whether temporal dynamics of the Pd differed between conditions. For this purpose, we analyzed the area under the curve in the contralateral vs. ipsilateral difference waveform with a repeated measures ANOVA with within subjects’ factors Time window (early: 100 – 200 ms, late: 200 – 35=00 ms) and Block type (fixed-features, mixed -features). Here, we focused on area under the curve rather than mean amplitude as this method is less sensitive to potential latency differences between individuals (note that mean amplitude yielded the same pattern of results). This analysis yielded no reliable interaction (*F* (1, 23) = 0.002, *p* = 0.97, 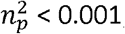; BFexcl = 3.70) suggesting that the observed modulation was uniform across time. This was also confirmed by an exploratory jackknife procedure (Miller, Patterson, & Ulrich, 1998) that did not identify a difference in onset latency between fixed- and mixed-feature conditions (thresh = 50% of maximum amplitude; onset_fixed_ = 140 ms, onset_mixed_ = 130 ms; *t* (23) = 0.08, *p* = 0.94).

The results thus far indicate that varying the target features (i.e., color and shape) randomly across trials uniformly attenuates the positivity elicited by lateral distractors in the typical Pd window. This raises the question to what extent the Pd in fixed-feature conditions as typically observed reflects sensitivity to target regularities across longer time scales above and beyond intertrial priming effects. Although our experiment was not designed to specifically target this question, in an exploratory analysis we aimed to address this by examining the effects of feature repetition in the mixed-feature condition. While behaviorally there was little to no difference between different forms of intertrial feature priming on the singleton presence benefit, in terms of the Pd the effect of intertrial color repetitions appeared most pronounced. For this purpose, we focused on the comparison of trials with and without repetition of the target color in the mixed-features condition. As visualized in Figure 3, the observed pattern of results was reminiscent of the main pattern of results comparing fixed- and mixed-feature conditions (see Figure 2), with no modulation of the N2pc, but an apparent attenuation of the Pd when the target color did not repeat from *one trial to the next. Indeed, whereas the N2pc elicited by lateral distractors was highly reliable irrespective of intertrial color priming (all t’s > 4*.*33, all p’s < 0*.*001, all d’s > 0*.*88), the distractor Pd reached significance on trials where target colors repeated (0*.*34* µ*V; t (23) = 2*.*51, p = 0*.*019, d = 0*.*51), but not on trials where the target color switched (0*.*16* µ*V; t (23) = 0*.*98, p = 0*.*34, d = 0*.*20). Note however that this pattern of results should be interpreted cautiously as the Prime (color repeat, color switch) by Hemifield interaction did not reach significance (F (1, 23) = 0*.*60, p = 0*.*45, = 0*.*025; BFexcl = 1*.*95) and future research is thus necessary to establish whether target color repetitions are by itself sufficient to modulate the Pd in response to a static color singleton*.

**Figure 3.**
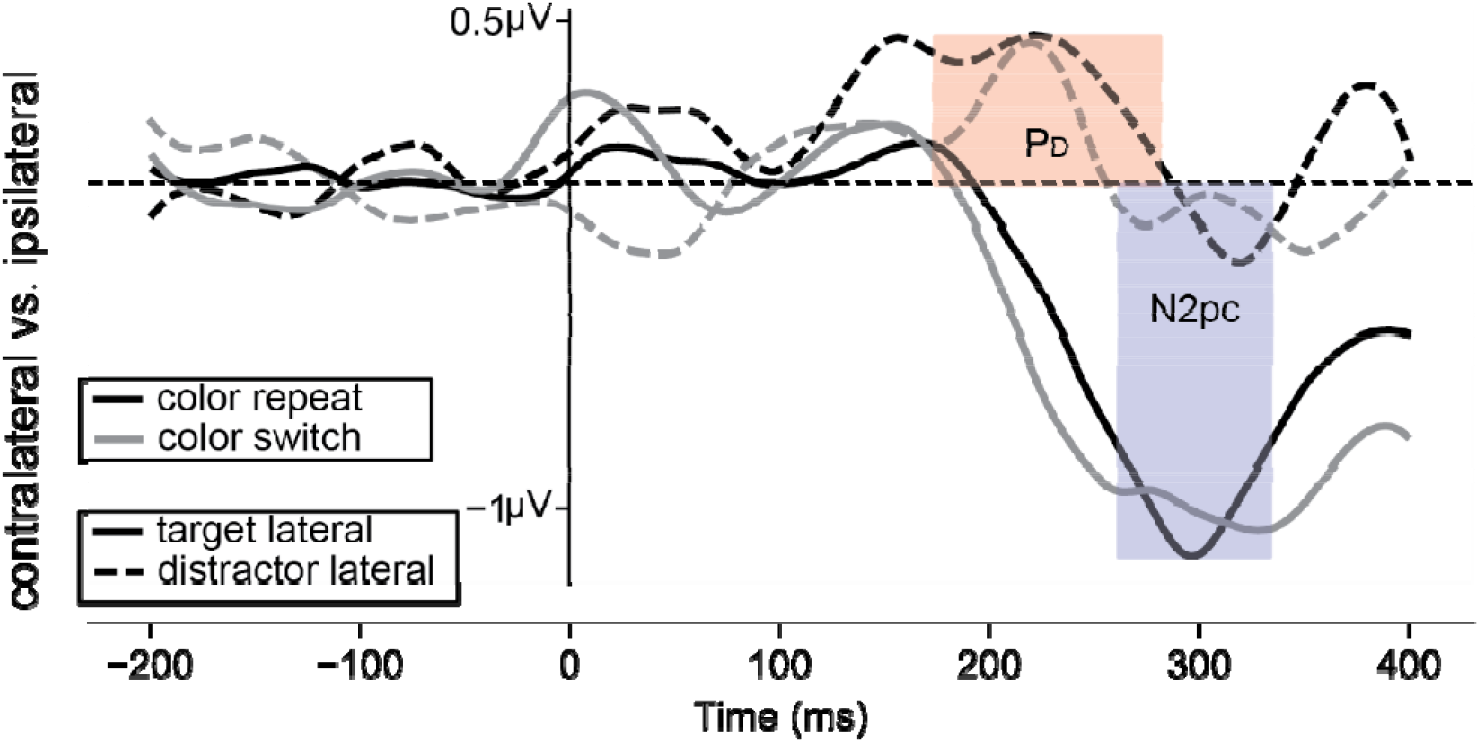
Target color repetitions appear to modulate the distractor Pd. Waveforms depict difference between contra- and ipsilateral waveforms at PO7/8 for target tuned (solid lines) and distractor tuned analyses (dashed lines), separately for color repeat (black) and color switch (grey) displays within the mixed-feature condition. Shaded areas reflect the time windows of interest for the N2pc (blue) and the Pd (red) analyses). The N2pc elicited by lateral targets did not differ between color repeat and color switch display (*m* _repeat_ = −1.0µV; *m* _switch_ = −1.0µV; ∆*m = 0*.*0*.; *n = 24;* two-tailed *p* = 0.95; *d* = 0.014; 95% CI = −0.41 – 0.39). By contrast, the Pd elicited by distractors appeared to be reliably attenuated when the target color switched although this was not supported by a reliable difference between mean amplitudes in the Pd window (*m* _repeat_ = 0.34µV; *m* _switch_ = 0.16µV; ∆*m = 0*.*18*.; *n = 24;* two-tailed *p* = 0.45; *d* = 0.16; 95% CI = −0.25 – 0.56). The waveforms in this plot were smoothed by a third-order polynomial (window length = 51) to improve the visibility of the effects but were analyzed using unsmoothed waveforms.

## General discussion

*The aim of the present study was to better characterize the P*_D_, *which has become one of the major tools to study suppression of salient distractors. Specifically, we aimed to examine the extent to which the P*_D_ *is driven by task-relevant regularities in designs typically used to study proactive distractor suppression (Chang & Egeth, 2019; Gaspelin & Luck, 2018a;* *Stilwell et al*., *2022**). To this end, we modified a heterogenous shape version of the* additional singleton paradigm (Theeuwes, 1991, 1992), which has been shown to reliably produce both behavioral and electrophysiological signatures of distractor suppression (Gaspelin & Luck, 2018a; Stilwell et al., 2022), such that target features were static in one half of the experiment (i.e., fixed-features condition), while they varied randomly in the other half of the experiment (i.e., mixed-features condition). This design made it possible to examine how processing of a fixed distractor color in a display of heterogenous shapes is shaped by expectations regarding the upcoming target features. We found that target selection was equally effective in both conditions, both in terms of manual response times, as well as in terms of the N2pc components. In contrast, while both conditions resulted in a singleton-presence benefits, the P_D_ elicited by lateralized distractors was reliably attenuated when the target features varied randomly across trials. This result demonstrates that the P_D_ elicited by distractors cannot unequivocally be attributed to a suppressive process, as it can also, at least when the experimental design allows, reflect upweighting of target features.

The present findings have important implications for the attentional capture debate, and specifically for studies that examine proactive distractor suppression via the P_D_ component. As noted in the introduction, below baseline suppression, a defining marker of proactive suppression, to date has been selective to experimental designs using heterogeneous shapes in which not only the distractor color, but also the defining target features (i.e., shape and color) are fixed across trials. Although it is notoriously difficult to empirically distinguish between feature specific upweighting and suppression (Gaspelin & Luck, 2018b), the consensus, as also acknowledged by recent formulations of the signal suppression hypothesis (Luck et al., 2021), is that under such conditions both upweighting and suppression concurrently guide attention (Chang & Egeth, 2019, 2021; Hamblin-Frohman, Chang, Egeth, & Becker, 2022; Vatterott & Vecera, 2012). Here, for the first time we demonstrate an electrophysiological correlate of this interplay between suppression an upweighting, by showing that the P_D_ is attenuated when target features are no longer static, but instead vary randomly across trials. This attenuation of the P_D_ under conditions that discourage target feature upweighting is consistent with a hybrid model of distractor processing, in which distractor inhibition is driven by two independent mechanisms. At the same time, it should be noted that it has even been argued that observed distractor inhibition can be exclusively explained by target feature upweighting (Oxner et al., 2022; Saenz, Buracas, & Boynton, 2002). In the current experimental design, when target features varied, they only did so between two possible options, leaving open the possibility that the remaining part of the P_D_ was still driven by target feature upweighting, but simply less pronounced upweighting given that in mixed-feature conditions it now had to be distributed across color space.

Although our results are consistent with the idea that the P_D_ elicited by distractor in heterogeneous search displays with static target features is sensitive to task relevant regularities, the exact underlying mechanism remains elusive. As outlined above, one possibility is that the P_D_ reflects both enhancement and suppression, such that the amplitude increases when the task also allows for secondary inhibition resulting from an upweighting of predictive target features (Noonan, Crittenden, Jensen, & Stokes, 2018; van Moorselaar & Slagter, 2020). Indeed, there is evidence that target representations can be strategically shifted off-veridical to optimally distinguish targets from distractors (Geng & Witkowski, 2019). Alternatively, it could be argued that the P_D_ purely reflects suppression, if one assumes that task relevant regularities influence how the distractor is encoded in relation to the other items in display (Becker, 2010). Whereas in the mixed-feature condition observers cannot rely on a relational distractor code and hence must rely on an absolute code, in the fixed-feature condition relational coding is possible to potentially strengthen distractor suppression. We believe this less likely however, given the behavioral evidence in support of target enhancement driving distractor inhibition in the current paradigm (Chang & Egeth, 2019, 2021; Oxner et al., 2022). Irrespective of the underlying mechanism however, the current results highlight the importance of taking target regularities into consideration when examining distractor suppression via the P_D._

The observation that the electrophysiological response elicited by salient distractors is modulated by target predictability is also consistent with previous ERP studies relying on homogeneous instead of heterogeneous search displays. When the target is defined as the unique shape in the display (e.g., a diamond among circles, or vice versa) there is some disagreement on the presence of a distractor P_D_ when the target shape varies unpredictable across trials, with some studies actually reporting an N2pc (Burra & Kerzel, 2013; Hickey, McDonald, & Theeuwes, 2006; Wang et al., 2019) suggesting attentional capture rather than distractor suppression (but see) (McDonald et al., 2013; van Moorselaar et al., 2021). By contrast, when the unique target shape is fixed, favoring feature search mode (Bacon & Egeth, 1994), not only the target N2pc increases, but distractors also reliably elicit a P_D_ (Burra & Kerzel, 2013; van Moorselaar et al., 2021). It should be noted however that even though a P_D_ signals suppression of distractors (Forschack, Gundlach, Hillyard, & Müller, 2022), in these displays distractors typically continue to interfere with attentional selection. To resolve this apparent discrepancy, one could argue that suppression was in place, but insufficiently so to counteract bottom-up attentional capture. Alternatively, the P_D_ may not necessarily signal suppression of a salient stimulus, but instead processing of a salient, yet irrelevant feature that does not require a further read out. Although highly speculative, such a framework would predict that any salient distractor would generate a P_D_, as long as the distractor does not resemble the target. In these circumstances, the initial capture by the salient distractor, does not require the formation of an object representation of the distractor as it can be immediately discarded as “not being the target”. Indeed, there is evidence that attention can be disengaged very rapidly if the distractor does not resemble the target (Born, Kerzel, & Theeuwes, 2011; Mulckhuyse, Van der Stigchel, & Theeuwes, 2009). In this framework, the P_D_ can still be envisioned as the mirror image of the N2pc, but not because they reflect covert shifts of attention (Eimer, 1996) or suppression (Gaspelin & Luck, 2018b), but instead indexing object individuation (Foster, Bsales, & Awh, 2020; Mazza & Caramazza, 2015) on the one hand in case of an N2pc and, and ignoring of specific features on the other hand in case of the P_D_.

An important caveat in the interpretation of our results is that targets and neutral non-targets always shared the same color and hence their effects cannot be disambiguated. Therefore, rather than attributing the observed P_D_ attenuation to an upweighting of static target features, an alternative explanation is to assume that the neural adaptation to repeated colors over time results in a reduced the inter-item competition between non-target items (Adam & Serences, 2021; Solomon & Kohn, 2014). According to such a framework however, the singleton should become increasingly less salient over time in the fixed-feature condition, which appears at odds with the observation that behaviorally the effect did not differ between conditions, and the fact that P_D_ decreased rather than increased in the mixed-feature condition where neural adaptation across trials should be less pronounced. Nevertheless, future work examining to what extent the P_D_ is sensitive to task relevant regularities should also take non-target regularities into account and also consider that the early positivity that we chose to label P_D_ in the current study, overlaps with the Ppc component which is proposed to be sensitive to feature discontinuities within search displays (Fortier-Gauthier, Moffat, Dell’Acqua, McDonald, & Jolicœur, 2012; Gokce, Geyer, Finke, Müller, & Töllner, 2014).

Our behavioral results indicate a finding what has been considered to be “a striking reversal of the capture effect” (Chang & Egeth, 2019) as observers were significantly faster on distractor present trials than on absent trials. This reversal of the capture effect has now been reported in several recent studies (Gaspelin et al., 2015; Lien, Ruthruff, & Hauck, 2022; Ma & Abrams, 2022; Stilwell & Gaspelin, 2021). While some may consider this reversal as surprising, this finding is consistent with our claim that during feature search in which the target is typically non-salient, the attentional window is adjusted to keep the discriminability of the target to an acceptable signal-to-noise-ratio (Liesefeld & Müller, 2020; Theeuwes, 2004, 2010). Because of the small attentional window, search proceeds serially and during serial search, there is no attentional capture by the salient distractor singleton. Because they are in serial search, participants can immediately discard the distractor which gives them one less item to inspect in distractor present than in absent trials (see Theeuwes, 2010, in press for a discussion). Consistent with this notion is our speculation that in these circumstances, in which search is serial, the distractor will generate a P_D_ because it can be immediately discarded as irrelevant and “not being the target”

In summary, the current study clearly demonstrates that in displays that encourage feature search mode, the P_D_ cannot unequivocally be attributed to suppression of distractor features, as at least in part, it is also sensitive to target regularities. This finding has large implications for future studies that use the P_D_ to examine whether specific distractor features can be proactively suppressed.

## Acknowledgements

This research was supported by a European Research Council (ERC) advanced grant (833029) to J.T. D.v.M designed the study, performed the analyses, and contributed most of the writing. C.H. collected the data and contributed to the writing. J.T. was closely involved in the design of the experiment and made significant contributions to the writing.

## Supplementary material

**Figure S1.**
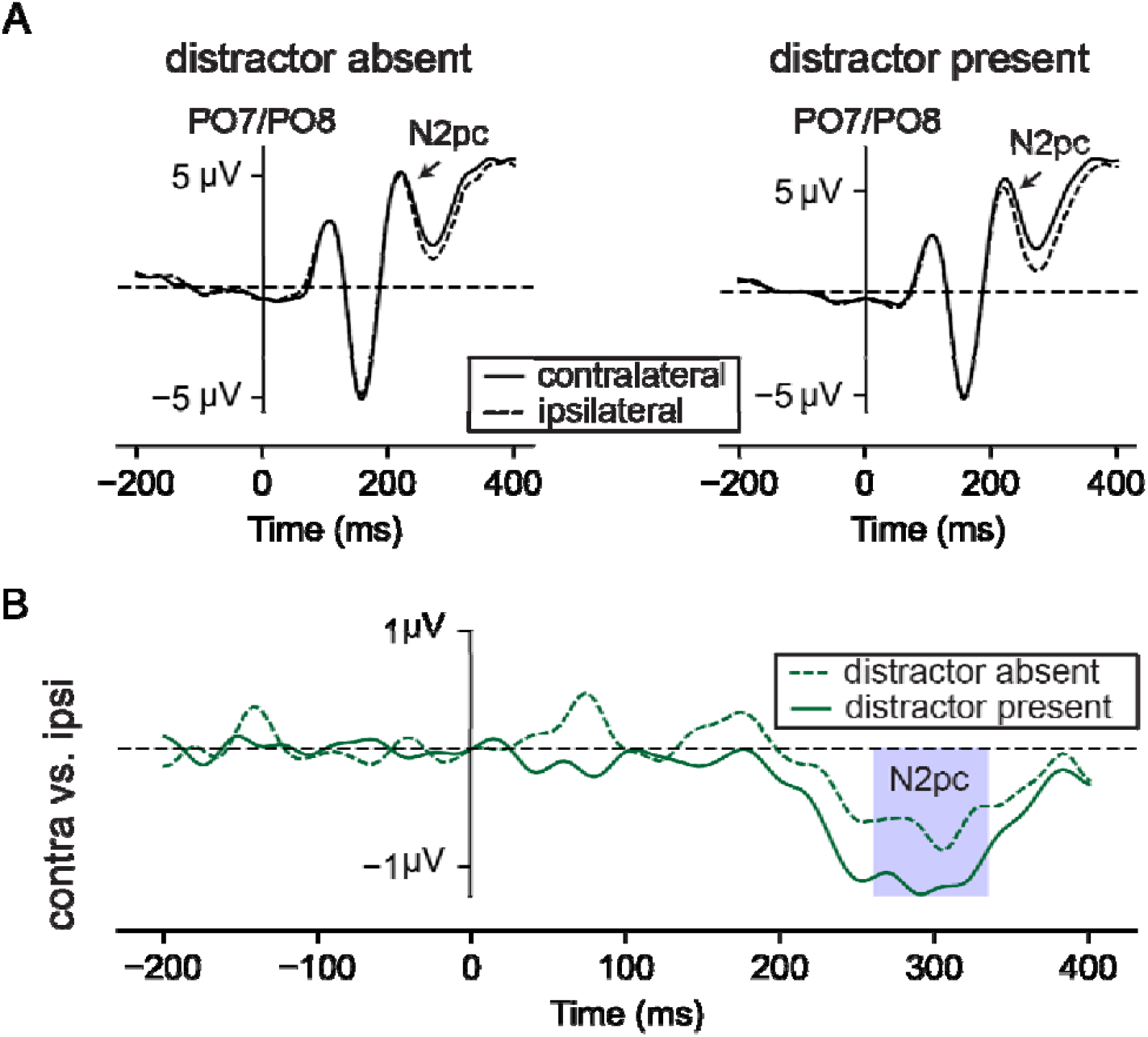
Target elicited waveforms in distractor absent and distractor present displays in the fixed-features condition. **A)** Electrophysiological results from search trials with lateral targets without a distractor (left) and with lateral targets accompanied by a distractor on the vertical midline (right). Ipsilateral (dashed lines) and contralateral (solid lines) waveforms reflect activity at electrode sides PO7/8. **B)** Difference waveforms between contra- and ipsilateral waveforms for target tuned waveforms on distractor absent (solid lines) and distractor present displays (dashed lines). The shaded area reflects the time windows of interest for the N2pc analyses). Despite the apparent numerical difference, the N2pc elicited by lateral targets did not differ between distractor absent and distractor present displays (*m* _absent_ = −0.54µV; *m* _present_ = −1.0µV; ∆*m = −0*.*5*.; *n = 24;* two-tailed *p* = 0.12; *d* = 0.33; 95% CI = −1.14 – 0.14).

**Figure S2.**
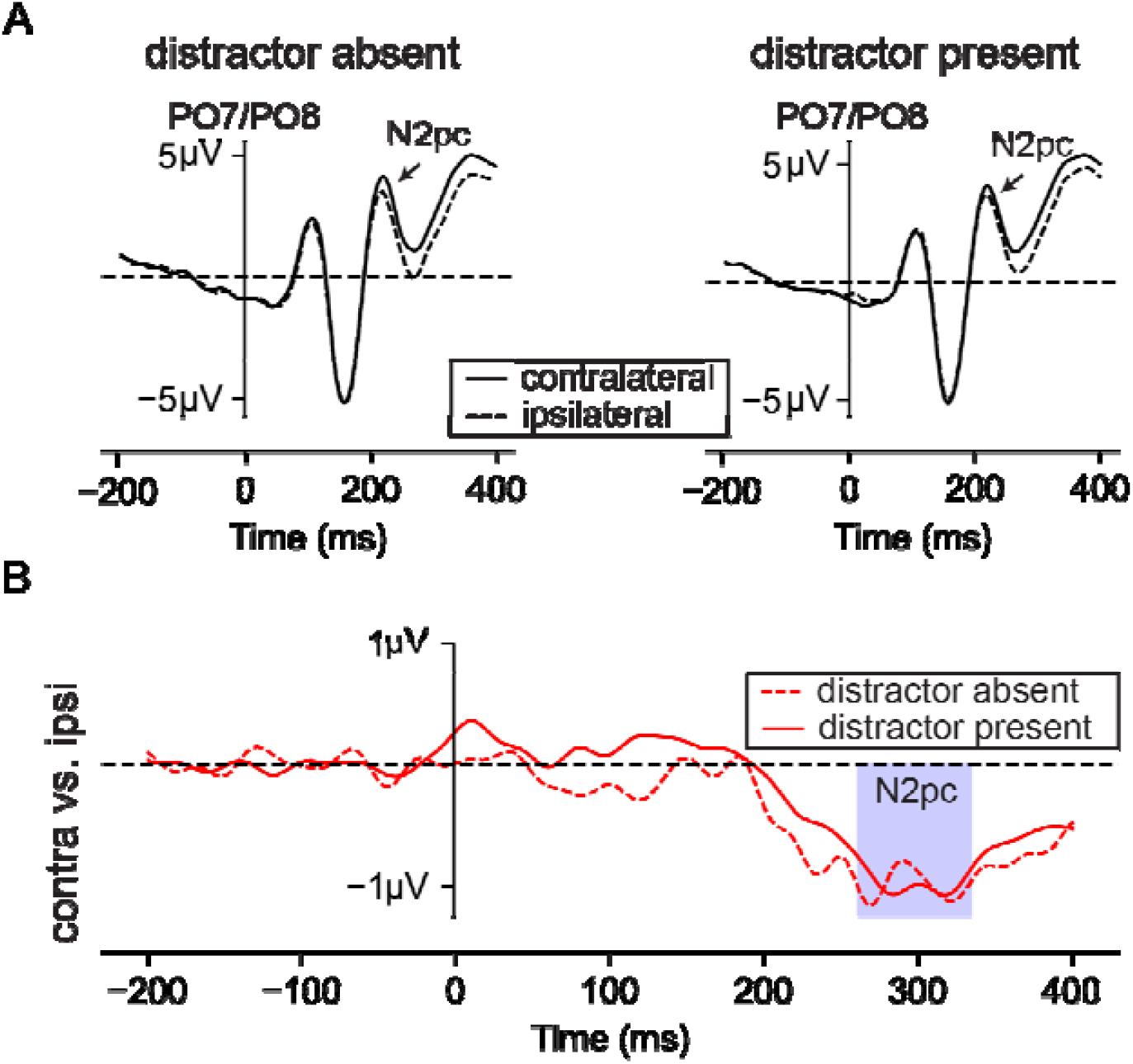
Target elicited waveforms in distractor absent and distractor present displays in the mixed-features condition. **A)** Electrophysiological results from search trials with lateral targets without a distractor (left) and with lateral targets accompanied by a distractor on the vertical midline (right). Ipsilateral (dashed lines) and contralateral (solid lines) waveforms reflect activity at electrode sides PO7/8. **B)** Difference waveforms between contra- and ipsilateral waveforms for target tuned waveforms on distractor absent (solid lines) and distractor present displays (dashed lines). The shaded area reflects the time windows of interest for the N2pc analyses). The N2pc elicited by lateral targets did not differ between distractor absent and distractor present displays (*m* _absent_ = −0.90µV; *m* _present_ = −0.77µV; ∆*m = −0*.*13*.; *n = 24;* two-tailed *p* = 0.52; *d* = 0.14; 95% CI = −0.54 – 0.28).

## Footnotes

1 The same pattern of results was obtained when time windows of interest were centered around condition-specific positive peaks in distractor tuned waveforms (177 – 287-ms and 171 – 282-ms for fixed- and mixed-features conditions, respectively), or alternatively when rather than a data driven approach time-windows matched the Pd window (i.e., 115 – 225-ms) reported in Stilwell et al. (2022).

